# Characterization of splice isoform switching during human kidney development

**DOI:** 10.1101/816371

**Authors:** Yishay Wineberg, Itamar Kanter, Nissim Ben-Haim, Naomi Pode-Shakked, Efrat Bucris, Tali Hana Bar-Lev, Sarit Oriel, Yishai Yehuda, Rotem Gershon, Rachel Shukrun, Dekel Dov Bar-Lev, Achia Urbach, Benjamin Dekel, Tomer Kalisky

**Affiliations:** Department of Bioengineering and Bar-Ilan Institute of Nanotechnology and Advanced Materials (BINA), Bar-Ilan University, Ramat Gan, Israel 52900; Pediatric Stem Cell Research Institute, Edmond and Lily Safra Children’s Hospital, Sheba Medical Center, Tel-Hashomer, Israel 52621; Division of Pediatric Nephrology, Sheba Medical Center, Tel-Hashomer, Israel 52621; Sackler Faculty of Medicine, Tel-Aviv University, Tel-Aviv, Israel 69978; The Mina and Everard Goodman Faculty of Life Sciences, Bar-Ilan University, Ramat-Gan, Israel 52900

**Keywords:** Kidney development, RNA sequencing, Alternative splicing, Mesenchymal to epithelial transition (MET), Wilms’ tumor

## Abstract

Nephrons are the functional units of the kidney. During kidney development, cells from the cap mesenchyme – a transient kidney-specific progenitor state – undergo a mesenchymal to epithelial transition (MET) and subsequently differentiate into the various epithelial cell types that create the tubular structures of the nephron. Faults in this transition can lead to a pediatric malignancy of the kidney called Wilms’ tumor that mimics normal kidney development. While kidney development has been characterized at the gene expression level, a comprehensive characterization of alternative splicing is lacking. We therefore performed RNA sequencing on cell populations representing early, intermediate, and late developmental stages of the human fetal kidney, as well as three blastemal-predominant Wilms’ tumor patient-derived xenografts. We identified a set of transcripts that are alternatively spliced between the different developmental stages. Moreover, we found that cells from the earliest developmental stage have a mesenchymal splice-isoform profile that is similar to that of blastemal-predominant Wilms’ tumors. RNA binding motif enrichment analysis suggests that the mRNA binding proteins ESRP1, ESRP2, RBFOX2, and QKI regulate mRNA splice isoform switching during human kidney development. These findings illuminate new molecular mechanisms involved in kidney development and pediatric kidney tumors.

**HIGHLIGHTS:**

- During fetal kidney development, kidney progenitor cells undergo a mesenchymal to epithelial transition (MET) and subsequently differentiate into the various epithelial cell types that create the tubular structures of the nephron.
- RNA sequencing identifies a set of transcripts that undergo splice isoform switching during the mesenchymal to epithelial transition (MET) that occurs in the course of human fetal kidney development.
- Cells in the early stages of kidney development have a mesenchymal splice-isoform profile that is similar to that observed in blastemal-predominant Wilms’ tumor patient-derived xenografts (WT-PDX) that represent an aggressive subtype of Wilms’ tumors.
- RNA binding motif enrichment analysis indicates that the mRNA binding proteins ESRP1, ESRP2, RBFOX2, and QKI regulate splice isoform switching during human kidney development.

## INTRODUCTION

Kidney development occurs during the embryonic stage from week 5 to week 36 of gestation in humans [1–4]. It starts as an interaction between two lineages that originate from the intermediate mesoderm: the ureteric duct – an epithelial tubular structure, and the metanephric mesenchyme – which is composed of loosely connected mesenchymal cells. This interaction causes the ureteric duct to invade the metanephric mesenchyme and, through a series of bifurcations, create a tree-like structure. The tips of this tree induce the surrounding cells of the metanephric mesenchyme to condense and form the “cap mesenchyme”, which is the nephric progenitor cell (NPC) population. Cells of the cap mesenchyme then undergo a mesenchymal to epithelial transition (MET) to create early epithelial structures called pre-tubular aggregates. These, in turn, progressively differentiate and elongate through a series of intermediate structures (renal vesicles, comma‐shaped, and S‐shaped bodies) to eventually give rise to all segments of the mature nephron.

Wilms’ tumor (WT) is the most common pediatric tumor of the kidney, with 75% of cases diagnosed in children under the age of five [1,5,6]. It is thought that Wilms’ tumors originate from fetal developing tissues that failed to differentiate properly. As a result, Wilms’ tumor is considered a model system for understanding the link between normal development and tumorigenesis. In many cases Wilms’ tumors contain three cellular components that appear in varying proportions between different patients and that mimic the three main stages of normal nephrogenic differentiation: the stroma – corresponding to the un-induced metanephric mesenchyme, the blastema - corresponding to the cap mesenchyme, and disordered non-functional epithelia that resemble the early epithelial structures of the fetal kidney. The blastema is thought to represent the least differentiated and most malignant component of the tumor and, typically, Wilms’ tumors with blastemal-predominance (after preoperative chemotherapy) require more aggressive treatment [7].

In a recent study [8] we found that cells from different developmental stages can be isolated from cultured human fetal kidney (hFK) cells using a combination of two surface markers: NCAM1, a cell adhesion protein that is overexpressed in the cap mesenchyme and early epithelial structures, and CD133 (PROM1), a membrane-bound protein which marks the more mature epithelial structures. A combination of those two markers allows for isolating cells from progressive stages of human fetal kidney development: the NCAM1+/CD133− cell fraction (which we denote “hFK1”) enriches for cells in the early renal developmental stages, mainly the cap mesenchyme; the NCAM1+/CD133+ cell fraction (“hFK2”) enriches for cells in an intermediate state corresponding to early renal epithelial structures; and the NCAM1–/CD133+ cell fraction (“hFK3”) enriches for cells at a more advanced differentiation state of the renal tubular epithelium.

To characterize these three cell fractions, we previously [8] performed RNA sequencing and qPCR. In hFK1 (the NCAM1+/CD133− cell fraction) we observed a strong expression of genes that were previously found to be over-expressed in the cap mesenchyme and in the un-induced metanephric mesenchyme (e.g. SIX2, SALL2, OSR1, and CDH11). Likewise, in hFK2 (the NCAM1+/CD133+ cell fraction) we observed an upregulation of genes related to epithelial differentiation (e.g. CDH1 and EPCAM), and even more in hFK3 (the NCAM1−/CD133+ cell fraction). This gradual pattern of switching from mesenchymal to epithelial characteristics, was further supported by splice isoform switching in the genes ENAH, CD44, and CTNND1, from their typical mesenchymal to epithelial isoforms, similar to what was previously observed in epithelial-mesenchymal transition (EMT) in embryonic development and metastatic breast cancer [9–13]. In a subsequent study [14] we used single-cell qPCR to confirm the splice isoform switching in the gene ENAH at the single-cell level. Moreover, we observed a strong over-expression of the mesenchymal isoform of ENAH in a blastemal-predominant Wilms’ tumor patient-derived xenograft (WT-PDX).

Although in those previous studies [8,14] we were able to find alternative splicing in specific genes, the RNA sequencing that we performed was originally designed for gene expression (50 bp, single-end, at approximately 20 million reads per sample) and was less suitable for comprehensive analysis of alternative splicing. We, therefore, present here a more thorough and quantitative study of the alternative splicing taking place in the developing human fetal kidney and in Wilms’ tumors. To achieve this, we re-sequenced the three cell fractions of the human fetal kidney (hFK1, hFK2, and hFK3) paired-end (2 × 126 bases) and more deeply (at approximately 40 million paired-end reads per sample). In addition, we also sequenced three additional samples of blastemal-predominant Wilms’ tumor xenografts that were derived from three different patients (WT11, WT14, and WT37) [8,15–18] (Fig. 1).

**Figure 1:**
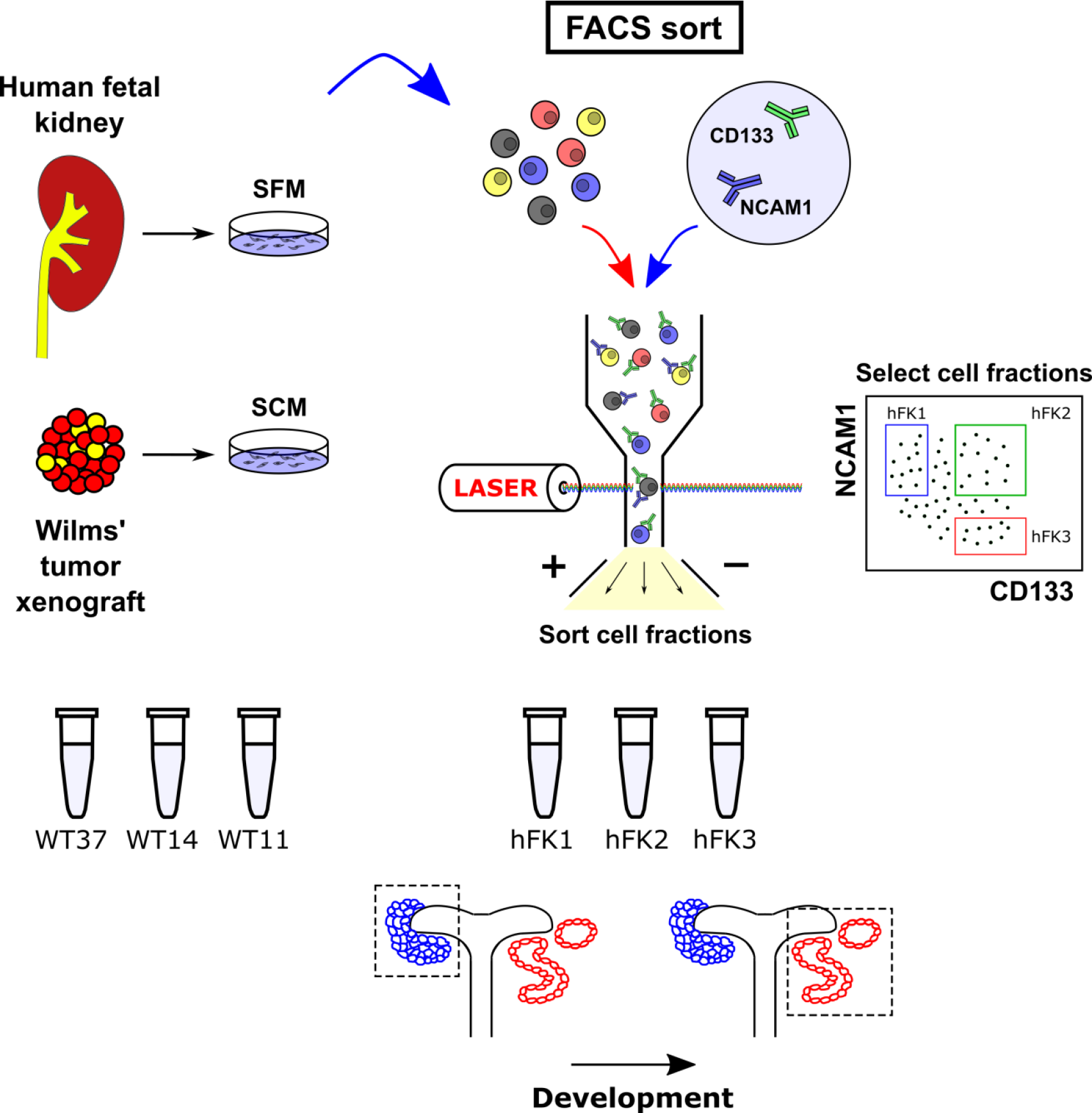
A sketch of the experiments performed to obtain the RNA samples that were analyzed in this study (see also [8]). Cells from a human fetal kidney (hFK) were cultured overnight in serum-free medium (SFM) which was found to help preserve a small population of undifferentiated cells. The cells were then dissociated and sorted by FACS to enrich for cell fractions representing three progressive renal developmental stages: NCAM-high/CD133-low (“hFK1”), a cell population that was previously found to be enriched for immature cells originating from the Cap mesenchyme; NCAM-high/CD133-high (“hFK2”), an intermediate state, presumably enriched for cells representing early epithelial structures; and NCAM-low/CD133-high (“hFK3”), a population enriched for more differentiated epithelial cells. In parallel, cells from three blastemal-predominant Wilms’ tumor patient-derived xenografts (“WT11”, “WT14”, and “WT37”) were cultured overnight in serum-containing medium (SCM). RNA from all samples was extracted and sequenced paired-end.

## RESULTS

### Cells at the early stage of human kidney development (NCAM-high/CD133-low, hFK1) have a mesenchymal gene expression profile that is similar to that observed in blastemal-predominant Wilms’ tumor patient-derived xenografts (WT-PDX)

We sequenced mRNA from three cell fractions from a human fetal kidney (hFK) that represent three consecutive renal developmental stages. The cells were cultured overnight in serum-free medium (SFM) which was found to help preserve a small population of undifferentiated cells, and then dissociated and sorted by FACS to enrich for the following cell fractions: NCAM-high/CD133-low (“hFK1”), a population that was previously shown to be enriched for immature cells originating from the cap mesenchyme; NCAM-high/CD133-high (“hFK2”), an intermediate state, presumably consisting of cells representing early developmental epithelial structures; and NCAM-low/CD133-high (“hFK3”), a population consisting of more differentiated epithelial cells. In parallel, cells from three blastemal-predominant Wilms’ tumor xenografts (WT-PDX) that were derived from three different patients (“WT11”, “WT14”, and “WT37”) were cultured overnight in serum-containing medium (SCM). RNA from all six samples was extracted, sequenced paired-end 2 × 126 bases at approximately 40 million paired-end reads per sample on an Illumina HiSeq 2500 platform, and analyzed for gene expression and alternative splicing.

For consistency, we first performed gene expression analysis for all samples. Principal Components Analysis (PCA) confirmed that the three fetal fractions (hFK1, hFK2, and hFK3) are sequentially ordered in gene expression space (Fig. 2A), as expected. The three Wilms’ tumor xenografts (WT11, WT14, and WT37) are closer to hFK1, the most immature fetal cell fraction, than to the other two fetal cell fractions (hFK2 and hFK3), which indicates that these Wilms’ tumors resemble an early stage in fetal kidney development. Likewise, we observed that genes associated with epithelial differentiation (PROM1 (CD133), EPCAM, and CDH1) increase towards hFK3, the cell fraction representing late-stage fetal development, whereas genes associated with the un-induced metanephric mesenchyme (ZEB1), the cap-mesenchyme (SIX2), or both (NCAM1 and CDH11) [14], decrease (Fig. 2A, red arrows). A heatmap of 67 genes that were previously found to be associated with the different cell types that co-exist within the developing kidney [14,19–21] (Fig. 2B), as well as barplots for selected genes associated with renal-mesenchymal and epithelial cell states (Fig. 2C), showed a sequential decrease in mesenchymal-associated genes and an increase in epithelial-associated genes in the fetal kidney samples (hFK1, hFK2, and hFK3), whereas all three Wilms’ tumor xenografts (WT11, WT14, and WT37) were observed to have a relatively high expression of mesenchymal associated genes.

**Figure 2:**
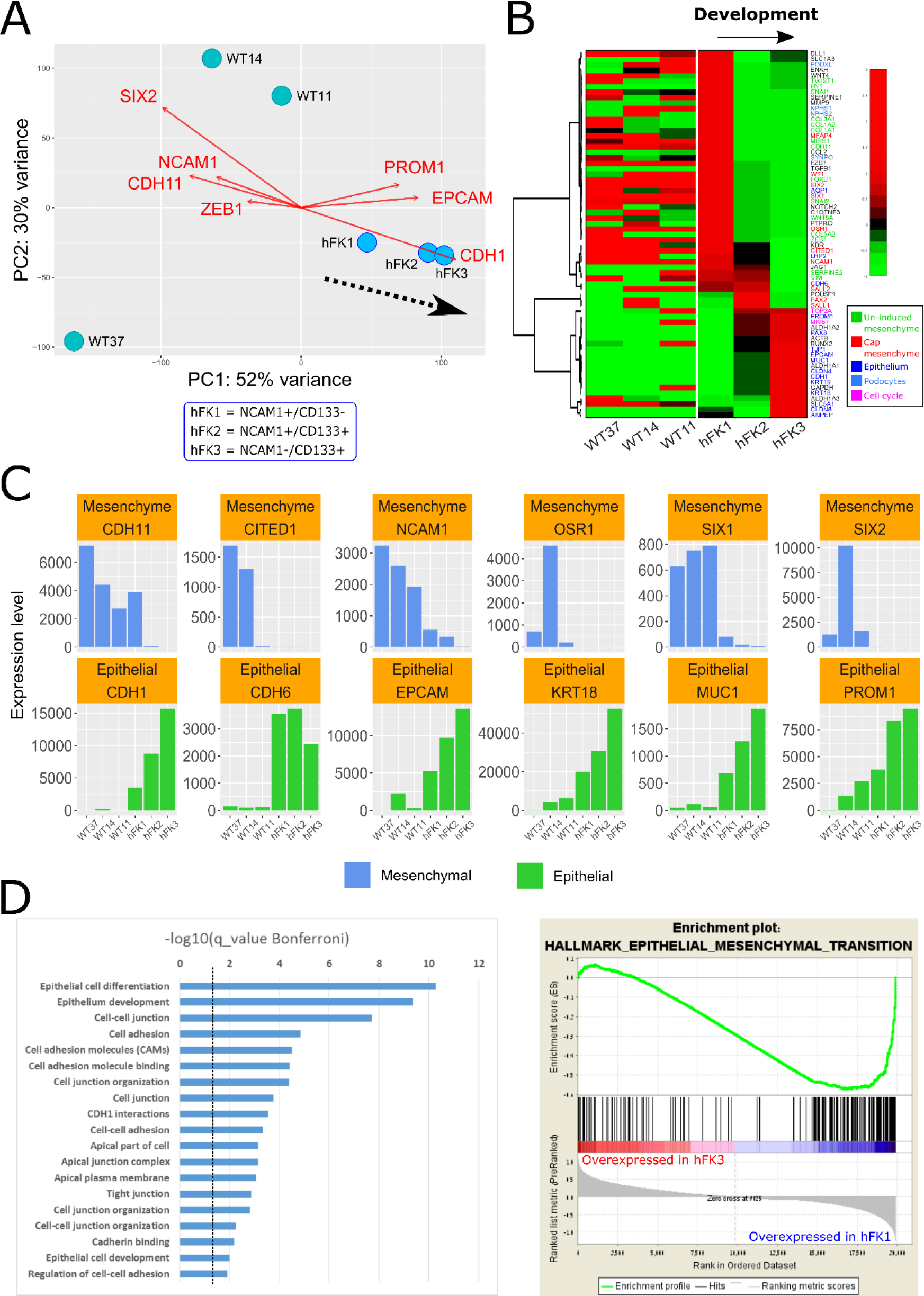
Cells at the early stage of human kidney development (NCAM-high/CD133-low, hFK1) have a mesenchymal gene expression profile that is similar to that observed in blastemal-predominant Wilms’ tumor patient-derived xenografts (WT-PDX). (A) A PCA biplot of gene expression levels. The three human fetal kidney cell fractions (hFK1, hFK2, and hFK3) lie on a trajectory (dotted black arrow) along which the epithelial-associated genes (CDH1, EPCAM, and PROM1 [=CD133]) increase and mesenchymal-associated genes (CDH11, ZEB1, NCAM1, SIX2) decrease. Note the large spread of Wilms’ tumor xenografts in gene expression space, which indicates a large variability between tumors from different patients. (B) Hierarchical clustering of 67 selected genes that were previously found to be related to kidney development. It can be seen that the blastemal-predominant Wilms’ tumor patient-derived xenografts (WT37, WT14, and WT11) are similar to hFK1 - the cell fraction that represents the most immature fraction of the human fetal kidney - in that they overexpress mesenchymal related genes and under-express epithelial related genes. The order of genes and the dendrogram were determined by hierarchical clustering of the human fetal kidney samples only (hFK1, hFK2, and hFK3). Note that although most epithelial associated genes that are over-expressed in hFK3, the podocyte markers PODXL, NPHS1/2, and SYNPO are only high in the early developmental stages (hFK1) and decrease with differentiation to hFK2 and hFK3. This is probably due to the fact podocytes cannot be cultured in the serum-free media that was used to culture the hFK cells. (C) Barplots of selected mesenchymal and epithelial associated genes involved in kidney development show sequential decrease in mesenchymal-associated genes and sequential increase in epithelial-associated genes and in the fetal kidney samples (hFK1, hFK2, and hFK3), whereas the three Wilms’ tumor xenografts (WT11, WT14, and WT37) all have high expression of mesenchymal associated genes and relatively low expression of epithelial-associated genes. (D) Gene Ontology (GO) enrichment analysis for 395 genes that were upregulated at least 2-fold (log2foldChange > 1) in hFK3 (the mature fetal developmental fraction) with respect to hFK1, WT11, WT14, and WT37 (see Fig. S1) shows that they are related to epithelial differentiation (see Table S2). Likewise, Gene Set Enrichment Analysis (GSEA) [23] showed that genes that are over-expressed in the early developmental cell fraction hFK1 (with respect to late fraction hFK3) are related to the Epithelial to Mesenchymal transition (EMT).

Next, we selected a set of 395 genes that were found by intersecting all the gene sets that were upregulated at least 2-fold (log2foldChange > 1) in hFK3 (the mature fetal developmental fraction) with respect to hFK1, WT11, WT14, and WT37 (see Fig. S1). We performed Gene Ontology (GO) enrichment analysis using ToppGene [22] and found that the genes in this set are related to epithelial differentiation (Fig. 2D, Table S2). Likewise, Gene Set Enrichment Analysis (GSEA) [23] of all genes, ranked according to their fold-change in expression between hFK1 and hFK3, showed that genes that are over-expressed in hFK1 (the early developmental cell fraction) vs. hFK3 (the late developmental cell fraction) are related to the Epithelial to Mesenchymal transition (EMT). All these findings indicate that cells at the early stage of kidney development (hFK1) have a mesenchymal gene expression profile that is similar to that of the blastemal-predominant Wilms’ tumor patient-derived xenografts (WT-PDX) (WT37, WT14, and WT11).

### Cells at the early stage of human kidney development (NCAM-high/CD133-low, hFK1) have a mesenchymal splice-isoform profile that is similar to that observed in blastemal-predominant Wilms’ tumor patient-derived xenografts (WT-PDX)

In order to check alternative splicing associated with the mesenchymal to epithelial transition (MET) that occurs during kidney development, we first inspected the genes ENAH [9–11], CD44 [12,13], CTNND1 [11,24], FGFR2 [24,25], and EPB41L5 [11,26] (Fig. S2), which were previously shown to be alternatively spliced in mesenchymal and epithelial tissues. We found that all three blastemal-predominant Wilms’ tumor patient-derived xenografts (WT37, WT14, and WT11) typically express the mesenchymal isoforms of these genes. In the fetal kidney, we found that hFK1 - the cell fraction representing the early stage of fetal kidney development - expresses a mixture of both mesenchymal and epithelial isoforms of these genes, whereas the cell fractions representing more differentiated stages (hFK2 and hFK3) predominantly express the epithelial splice isoforms.

We next performed a more comprehensive analysis using rMATS [27], a tool for detecting alternative splicing from RNA sequencing datasets. PCA of exon inclusion levels (Fig. 3C) showed that the three human fetal kidney samples (hFK1, hFK2, and hFK3) lie on a trajectory along which the epithelial-associated exons within the genes CD44 and ENAH sequentially increase and a mesenchymal-associated exon within the gene CTNND1 sequentially decreases. We also observed that the exon inclusion level profiles of the blastemal-predominant Wilms’ tumor patient-derived xenografts are typically closer to hFK1 - the most immature fetal cell fraction - than to the other more mature fetal cell fractions (hFK2 and hFK3), similar to what we have seen in gene expression (Fig. 2A).

**Figure 3:**
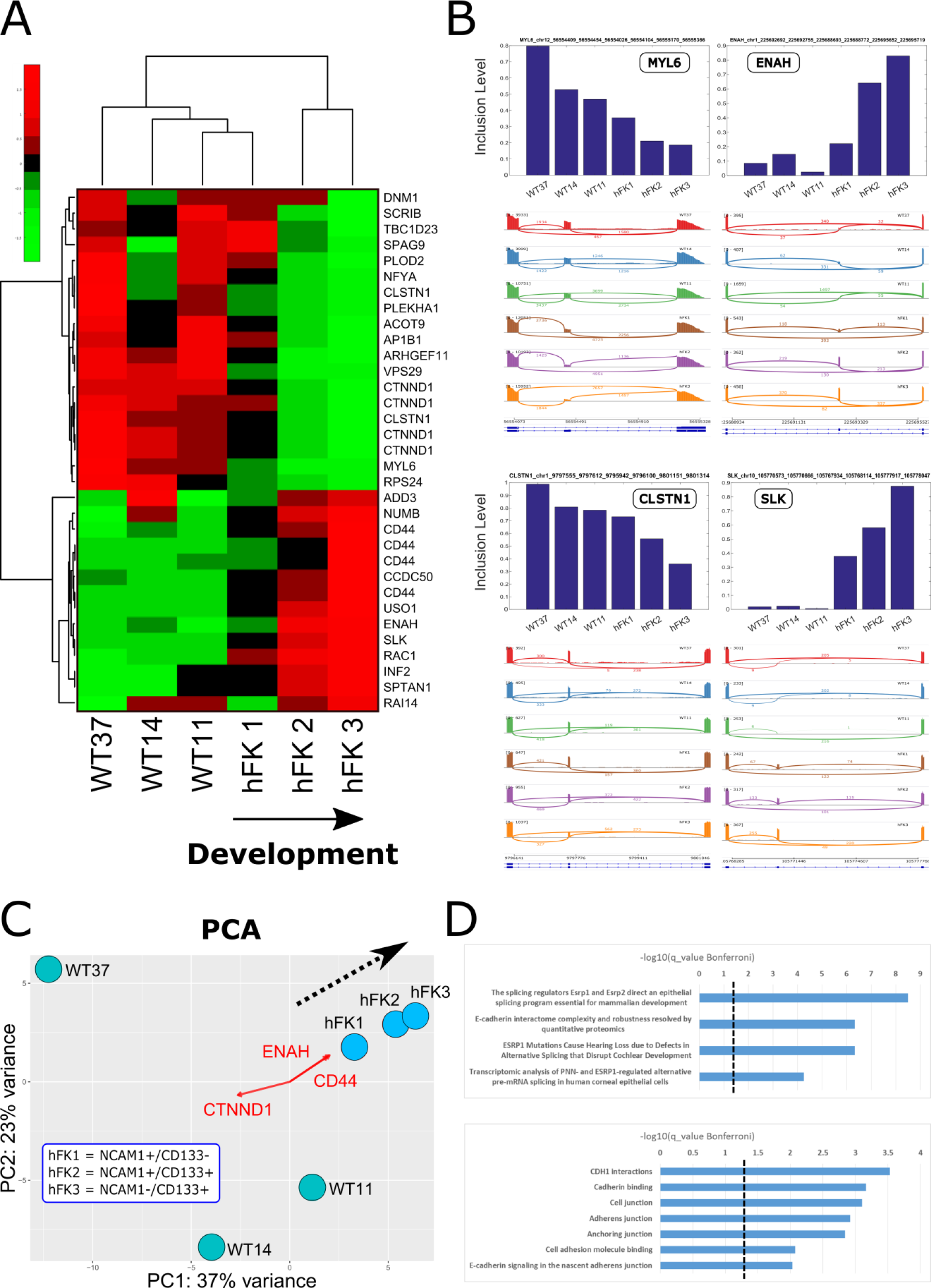
Cells at the early stage of human kidney development (NCAM-high/CD133-low, hFK1) have a mesenchymal splice-isoform profile that is similar to that observed in blastemal-predominant Wilms’ tumor patient-derived xenografts (WT-PDX). (A) Hierarchical clustering of the inclusion levels of 33 selected cassette exons that were manually found to be alternatively spliced between hFK1 and hFK3 - the two cell populations that represent early (hFK1) and late (hFK3) developmental stages in the fetal human kidney. Note that the late-stage fetal kidney cell fractions hFK2 and hFK3 are grouped in one cluster, whereas hFK1, the early-stage fraction, is grouped with the Wilms’ tumor xenograft samples. The 33 cassette exons were selected as follows: We chose cassette exons that were significantly differentially spliced (FDR < 1E-9 and difference in inclusion levels > 0.1) between hFK1 and hFK3, and from these we selected 33 cassette exons that also showed clear alternative splicing by manual inspection in the IGV genome browser. (B) Barplots and sashimi plots for selected cassette exons show the change in inclusion levels between the different cell fractions. Exons within the genes MYL6 and CLSTN1 are high in Wilms’ tumors and early fetal kidney cells (hFK1) and decrease during kidney development (hFK2 and hFK3), while those within ENAH and SLK are low in Wilms’ tumors and early fetal kidney cells (hFK1) and increase during kidney development (hFK2 and hFK3). (C) A PCA biplot of exon inclusion levels that were calculated by rMATS. Each point represents a different cell fraction. The three human fetal kidney samples (hFK1, hFK2, and hFK3) lie on a trajectory (dotted black arrow) along which the epithelial-associated exons within the genes CD44 and ENAH sequentially increase, and a mesenchymal-associated exon within the gene CTNND1 sequentially decreases. For PCA analysis we used all cassette exons that were detected by rMATS. (D) Gene Ontology (GO) enrichment analysis for the genes containing the 33 selected cassette exons indicates that they are related to epithelial differentiation and that alternative splicing in these genes is regulated by the splicing regulators ESRP1 and ESRP2 (see Table S4).

We next chose cassette exons that were significantly differentially spliced (FDR < 1E-9 and difference in inclusion levels > 0.1) between hFK1 and hFK3 - the fractions representing the earliest and latest stages of kidney development – and of these we selected 33 cassette exons that also showed clear alternative splicing by manual inspection in the IGV genome browser [28]. Hierarchical clustering (Fig. 3A) and barplots (Fig. 3B) of the inclusion levels of these 33 selected cassette exons showed that hFK1, the early-stage fetal fraction, has a splice isoform profile that is similar to the Wilms’ tumor xenografts (WT37, WT14, and WT11). Gene Ontology (GO) enrichment analysis for the genes containing these 33 selected cassette exons showed that they are related to epithelial differentiation (fig. 3D, Table S4). Moreover, enrichment analysis with respect to gene sets found in previous studies (e.g. [29]) indicated that alternative splicing in these genes is regulated by the splicing regulators ESRP1 and ESRP2 (fig. 3D, Table S4).

We note that we did not find any consistent relationship between gene expression levels and exon inclusion levels within the same gene (Fig. S3): For some genes we observed a very positive correlation (e.g. USO1, in which both expression levels and exon inclusion levels increase during development; r=0.83), while for others we observed a very negative correlation (e.g. ACOT9, in which expression levels increase during development and exon inclusion levels decrease; r=−0.77) or no correlation at all (e.g. SLK, in which expression levels are constant while and exon inclusion levels increase during development; r=0.086).

### RNA binding motif enrichment analysis indicates that the mRNA binding proteins ESRP1, ESRP2, RBFOX2, and QKI regulate splice isoform switching during human kidney development

Next, we searched among known RNA binding proteins (RBP’s) for genes that are likely to be responsible for regulating splice isoform switching during human kidney development [30,31]. Among the genes that we observed to be alternatively spliced are FAT1 [9,32] and PLOD2 [9,33,34] (Fig. S2), two genes for which alternative splicing was previously found to be regulated by the RNA binding protein RBFOX2 [9], and ARHGEF10L [29,35,36], a gene for which alternative splicing was previously found to be regulated by the RNA binding proteins ESRP1 and ESRP2 [29].

In order to conduct a more systematic search, we compared the mean expression levels of 89 known RNA binding proteins [37–39] and found several putative splicing regulators that were differentially expressed between hFK1, the cell fraction representing the early stages of fetal kidney development, and hFK3, the cell fraction representing the latest most differentiated stage (Fig. 4A). For example, we found that ESRP1 and ESRP2 are over-expressed in hFK3 (Fig. 4B, Fig. S5A), and RBFOX2 is over-expressed in hFK1 (Fig. 4B).

**Figure 4:**
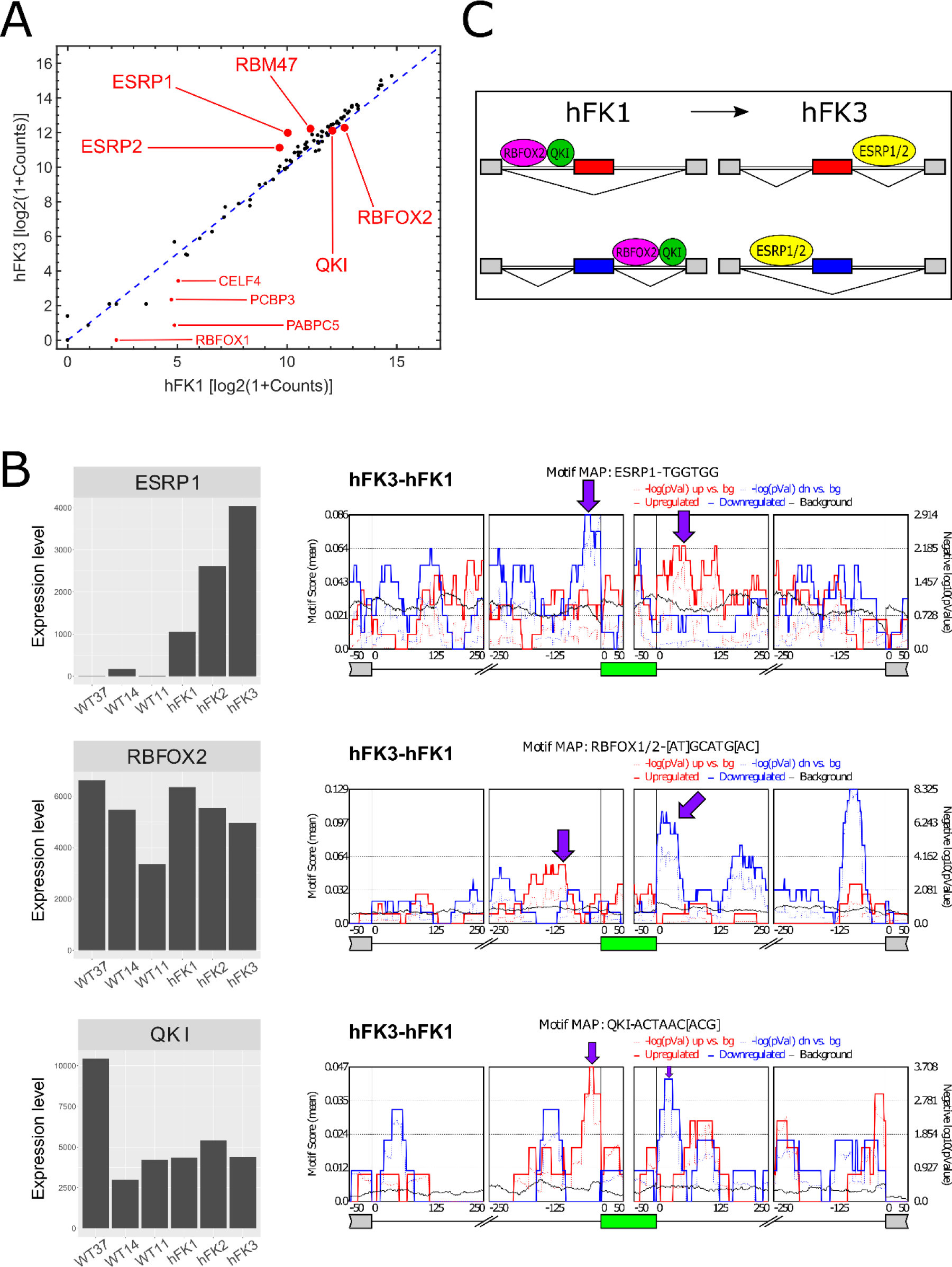
RNA binding motif enrichment analysis indicates that the mRNA binding proteins ESRP1, ESRP2, RBFOX2, and QKI regulate splice isoform switching during human kidney development. (A) Shown is a comparison of gene expression levels of 89 known RNA binding proteins between hFK1 and hFK3 (see Methods). (B) RNA binding motif enrichment analysis using rMAPS identifies four putative splicing factors: ESRP1, ESRP2, RBFOX2, and QKI. ESRP1 and ESRP2 (see also Fig. S5A) have low expression levels in all Wilms’ tumor samples and monotonically increase along kidney development, starting with moderate levels in hFK1 and reaching a maximum in hFK3. Likewise, exons that were enhanced in hFK3 (with respect to hFK1) are enriched for ESRP1 binding sites at their downstream 3’ flanking intron (red curve), while exons that are silenced in hFK3 are enriched for ESRP1 binding sites at their upstream 5’ flanking intron (blue curve). On the other hand, RBFOX2 shows a monotonic decrease in expression levels along kidney development, starting with high levels in hFK1 and decreasing in hFK2 and hFK3. Exons that are elevated in hFK1 (with respect to hFK3) are enriched for RBFOX2 binding sites at their downstream 3’ flanking intron (blue curve). Although QKI did not show an appreciable change in expression levels between hFK1, hFK2, and hFK3, its RNA binding sites show similar behavior to RBFOX2, that is, exons that are elevated in hFK1 (with respect to hFK3) are enriched for QKI binding sites at their downstream 3’ flanking intron (blue curve), and exons that are elevated in hFK3 (with respect to hFK1) are enriched for QKI binding sites at their upstream 5’ flanking intron (red curve). (C) These results are consistent with the model for splicing regulation during the Mesenchymal to Epithelial Transition (MET) as proposed by Yang et al [34]. Applying this model to kidney development, the hFK1 cell fraction corresponds to early kidney developmental stages and is predominantly composed of mesenchymal cells (the cap mesenchyme and the un-induced metanephric mesenchyme). Therefore in hFK1, ESRP1 and ESRP2 are low and RBFOX2 and QKI promote exon inclusion by binding to downstream introns, or exon skipping by binding to upstream introns. The hKF3 fraction corresponds to a late more differentiated kidney developmental stage and is predominantly composed of epithelial cells. As a result, ESRP1 and ESRP2 are high in hFK3 and promote exon inclusion by binding to downstream introns, or exon skipping by binding to upstream introns.

We next used rMAPS [29] to perform enrichment analysis for RNA binding sites (motifs) that belong to these known RNA binding proteins (RBP’s). We found that the RNA binding sites of ESRP1, ESRP2, and RBFOX2 are enriched in the upstream or downstream neighboring introns of the cassette exons that are differentially expressed between hFK1 and hFK3 (Fig. 4B, Fig. S5A) [29,34,36,38,40,41]. This indicates that ESRP1, ESRP2, and RBFOX2 are splicing regulators involved in the Mesenchymal to Epithelial Transition (MET) that occurs during human fetal kidney development (Fig. 4C). Another putative regulator is QKI1 which, although does not show an appreciable change in expression levels between hFK1 and hFK3, has an RNA binding site that shows a similar enrichment pattern as RBFOX2 [34].

These results are somewhat similar to what was previously observed by Yang *et al.* [34] in cells from a human H358 epithelial non-small cell lung cancer (NSCLC) cell line undergoing EMT (see comparison in Fig. S4) and consistent with their proposed model for splicing regulation during the Mesenchymal to Epithelial Transition (MET) (Fig. 4C). Applying this model to kidney development, the hFK1 cell fraction corresponds to early kidney developmental stages and is predominantly composed of mesenchymal cells (the cap mesenchyme and the un-induced metanephric mesenchyme). Therefore in hFK1, ESRP1 and ESRP2 are low and RBFOX2 and QKI promote exon inclusion by binding to downstream introns, or exon skipping by binding to upstream introns. The hKF3 fraction corresponds to a later more differentiated kidney developmental stage and is predominantly composed of epithelial cells. As a result, ESRP1 and ESRP2 are high in hFK3 and promote exon inclusion by binding to downstream introns, or exon skipping by binding to upstream introns.

## DISCUSSION

In this study, we used “bulk” RNA sequencing in order to comprehensively characterize alternative splicing in cell populations representing early, intermediate, and late developmental stages of the human fetal kidney, as well as three blastemal-predominant Wilms’ tumor patient-derived xenografts (WT-PDX) that represent an aggressive subtype of Wilms’ tumors. We found a set of transcripts that are alternatively spliced between the different developmental stages and identified putative splicing regulators. Moreover, we found that the blastemal-predominant Wilms’ tumor patient-derived xenografts (WT-PDX) resemble the earliest developmental stage in both gene expression and alternative splicing. These results illuminate new molecular mechanisms involved in kidney development and may assist in the design of new markers for pediatric kidney tumors.

The kidney is a heterogeneous organ composed of millions of microstructures, each composed of different cell types. Ideally, the best way to characterize alternative splicing throughout the different developmental stages is by applying full transcript length single-cell RNA sequencing [42–45]. We have recently performed full transcript length single-cell RNA sequencing in the developing mouse fetal kidney [46] and identified a set of transcripts - similar to those found here - that undergo splice isoform switching during the transition between mesenchymal and epithelial cellular states that takes place in the course of mouse fetal kidney development. Likewise, RNA binding motif enrichment analysis suggested that Esrp1/2 and Rbfox1/2 are splicing regulators of the Mesenchymal to Epithelial Transition (MET) that occurs during mouse kidney development, also similar to what we found here.

However, single-cell RNA sequencing is costly and often prone to bias and low coverage, which makes alternative splicing analysis challenging. Moreover, human fetal tissues are technically difficult to obtain in a “fresh” and viable state that is suitable for single-cell RNA sequencing. Therefore, for the present study, we deeply sequenced “bulk” RNA from cells that were harvested from primary human fetal tissues, cultured overnight, and enriched by flow cytometry with fluorescently labeled antibodies according to a protocol that we have previously shown to enrich for cell fractions that represent early, intermediate, and late developmental stages of the human fetal kidney [8]. Although some cells types such as podocytes are not preserved during the overnight culturing, we have previously shown [8] that this method does faithfully represent central developmental processes that occur during kidney development such as the Mesenchymal to Epithelial Transition (MET). Likewise, in order to overcome similar difficulties in obtaining “fresh” Wilms’ tumor samples, we used patient-derived xenografts (WT-PDX) that were established in the course of previous studies [8,15–18].

We note that RNA binding motif enrichment analysis is currently somewhat limited, probably due to limited knowledge and the fact that many known motifs are typically only a few bases long. This results in non-specific results pointing to candidate splicing regulators that are expressed at low levels in some populations (e.g., RBFOX1, Fig. 4A) or, conversely, in potential splicing regulators that are differentially expressed but for which no motif enrichment is detected. For example, RBM47 [34] is differentially over-expressed in hFK3 vs. hFK1 (Fig. 4A, Fig. S5B), which indicates that it might also be involved in splicing regulation during kidney development, but we did not find any significant motif enrichment for this gene. Likewise, we observed additional RNA binding proteins (Fig. S6) that are differentially expressed between hFK1 and hFK3 or between the Wilms’ tumors (WT37, WT14, and WT11) and hFK3, and these might also be involved in splicing regulation.

Although in this study we did not perform functional validation, the putative splicing regulators that we identified here (ESRP1, ESRP2, RBFOX2, and QKI) were found to have similar functionality in other developing organs and in-vitro systems [11,24,26,34,36,40,47]. Moreover, it was recently shown that Esrp1 ablation in mice, alone or together with Esrp2, results in reduced kidney size, fewer ureteric tips, reduced nephron numbers, and a global reduction of epithelial splice isoforms in the transcriptome of ureteric epithelial cells [48]. Our results indicate that ESRP1 and ESRP2 may play a similar role also in human kidney development. Moreover, the fact that kidneys still develop in mice after Esrp1 and Esrp2 ablation, taken with our results, suggests that other splicing regulators such as Rbfox1, Rbfox2, or Qki can compensate, although partially, for the ablation of Esrp1 and Esrp2.

## METHODS

### Tissue collection, dissociation, culturing, and flow cytometry

In this study we characterized alternative splicing in three cell populations from the human fetal kidney (“hFK1”, “hFK2”, and “hFK3”) that we have shown in a previous study [8] to represent progressive developmental stages of the human fetal kidney. In that study [8], we performed RNA sequencing that was designed for gene expression analysis (50 bp, single-end, at approximately 20 million reads per sample) and was less suitable for comprehensive analysis of alternative splicing. We, therefore, re-sequenced the three cell fractions paired-end (2 × 126 bases) and more deeply (at approximately 40 million paired-end reads per sample). In addition, we also sequenced three additional RNA samples of blastemal-predominant Wilms’ tumor xenografts (WT-PDX) that were derived from three different patients (WT11, WT14, and WT37). These RNA samples were collected in the course of previous studies [8,15–18] that were performed beforehand in the Pediatric Stem Cell Research Institute at the Sheba Medical Center. Details of tissue collection can be found in the above references, while below we describe them briefly.

Human fetal kidney cells were collected as previously described [8,15] (Fig. 1). Briefly, kidneys were collected from elective abortions at fetal gestational age between 15 to 19 weeks. The kidneys were dissociated into single-cell suspensions, resuspended in serum-free medium (SFM), plated in flasks, and cultured until reaching 80% confluence. FACS was used to isolate three cell fractions representing progressive developmental stages of the human fetal kidney: hFK1 (NCAM1+/CD133−), hFK2 (NCAM1+/CD133+), and hFK3 (NCAM1−/CD133+), as previously described [8]. All assays were conducted with low passage cultured cells (passage 0). Likewise, Wilms’ tumor patient-derived xenografts (WT-PDX) were established, propagated in mice, and collected as previously described [8,17] (Fig. 1). The xenografts were then dissociated into single-cell suspensions, resuspended in serum-containing medium (SCM), plated in flasks, and cultured overnight. For this study we used three blastemal-predominant Wilms’ tumor xenografts (WT-PDX) originating from three different patients (WT11, WT14, and WT37).

All human tissue handling procedures were approved by the local ethical committee of the Sheba Medical Center and informed consent was given by the legal guardians of the patients involved according to the Declaration of Helsinki. Likewise, all animal procedures were approved by the Institutional Animal Care and Use Committee in the Sheba Medical Center.

### RNA purification and sequencing

Bulk total RNA was prepared from ~1.5*10^5^ cells using the Direct-zol RNA MiniPrep kit (R2050, Zymo Research) according to the manufacturer’s instructions and stored in − 80 °C. RNA was quantified on an Agilent BioAnalyzer (Agilent Technologies) and aliquots of 270–500 ng were made into cDNA libraries using the TruSeq mRNA-Seq library kit (Illumina).

All 6 libraries (hFK1, hFK2, hFK3, WT11, WT14, and WT37) were sequenced paired-end 2 × 126 bases on an Illumina HiSeq 2500 platform in the Israel National Center for Personalized Medicine (G-INCPM). For each sample we obtained approximately 40 million paired-end reads.

### RNA sequence data preprocessing and gene expression analysis

Raw reads were aligned by TopHat2 [49] to the human hg19 genome. Aligned reads were counted by HTSeq [50]. Data normalization (resulting in a matrix of normalized gene expression counts), estimation of size factors, and differential gene expression were done by DESeq2 [51].

The GEO series record for the sequencing data is: [*TBD*]

Heatmaps, PCA biplots, and barplots were performed in Matlab (Mathworks) and R. Gene Ontology (GO) enrichment was done with ToppGene (https://toppgene.cchmc.org) [22]. Gene Set Enrichment Analysis (GSEA) [23] was used to check for enrichment of gene sets from the Molecular Signatures Database (MSigDB).

### Splicing analysis: Identification of splicing events and putative splicing regulators

rMATS [27] was used to detect splice isoform switching events between the cell fractions representing early (hFK1) and late (hFK3) stages of human fetal kidney development, as well as for quantifying inclusion levels in all transcriptomes. Selected splicing events (e.g. cassette exons) were visualized and validated using IGV [28] and Sashimi plots [52]. Gene Ontology (GO) enrichment of genes containing differential splicing events was done with ToppGene (https://toppgene.cchmc.org) [22].

rMAPS (http://rmaps.cecsresearch.org/) [37] was used to test for enrichment of binding motifs of RNA binding proteins (RBP’s) in the vicinity of alternatively spliced cassette exons in order to identify putative splicing regulators.

A list of 89 RNA binding proteins (RBP’s) was obtained from the rMAPS website (http://rmaps.cecsresearch.org/Help/RNABindingProtein) [37–39]. Apart from the RNA binding motifs that are tested by the default settings in the rMATS website, we also tested additional UGG-enriched motifs that were previously found to be binding sites for the RNA binding proteins ESRP1 [29,40] and ESRP2 [36] (Table S5). For the RNA binding proteins RBFOX1 and RBFOX2, following [34] and the CISBP-RNA database [38] (http://cisbp-rna.ccbr.utoronto.ca) we assumed that both proteins (RBFOX1 and RBFOX2) preferentially bind to the same motif ([AT]GCATG[AC]) on mRNA.

## AUTHOR CONTRIBUTIONS

Study initiation and conception – Y.W., I.K., N.B.H., E.B., N.P.S., B.D., and T.K.; Human Fetal kidney collection, culturing, sorting, and RNA extraction – N.P.S. and R.G.; Wilms’ tumor patient-derived xenograft (WT-PDX) maintenance, collection, culturing, and RNA extraction – N.P.S., R.G, R.S., and D.D.B.L.; RNA quality checks – E.B. and T.H.B.L.; RNA sequence preprocessing – I.K., N.B.H., and E.B.; Gene expression and splicing analysis – Y.W., I.K., E.B., N.B.H., and T.K.; Other intellectual contribution – S.O., Y.Y., and A.U.; Manuscript writing – Y.W., I.K., and T.K.

## ACKNOWLEDGEMENTS

We wish to thank Jordan Kreidberg, Steve Potter, Oded Volovelsky, Morris Nehama, Tal Shay, Rotem Karni, Peter Hohenstein, and all members of our labs for useful comments and suggestions.

## SUPPORTING INFORMATION LEGENDS

**Supplementary information:** Supplementary figures.

**Table S1:** Gene expression values.

**Table S2:** Gene Ontology (GO) enrichment analysis results from ToppGene for the set of 395 genes that were found by intersecting all the genes that were upregulated at least 2-fold (log2foldChange > 1) in hFK3 (the mature fetal developmental fraction) with respect to hFK1 (the immature fraction), WT11, WT14, and WT37 (see Figs. 2D and S1).

**Table S3:** rMATS tables of alternatively spliced cassette exons.

**Table S4:** Gene Ontology (GO) enrichment analysis results from ToppGene for the genes containing the 33 selected cassette exons (see also Fig. 3A). These 33 selected cassette exons were significantly differentially spliced (FDR < 1E-9 and difference in inclusion levels > 0.1) between hFK1 and hFK3 – the cell fractions representing the earliest and latest stages of kidney development - and were also found to show clear alternative splicing by manual inspection in the IGV genome browser.

**Table S5:** A list of RNA motifs used for identifying putative splicing regulators.

**Program:** A compressed directory containing programs and datasets for data visualization.

## Supplementary figures

**Figure S1:**
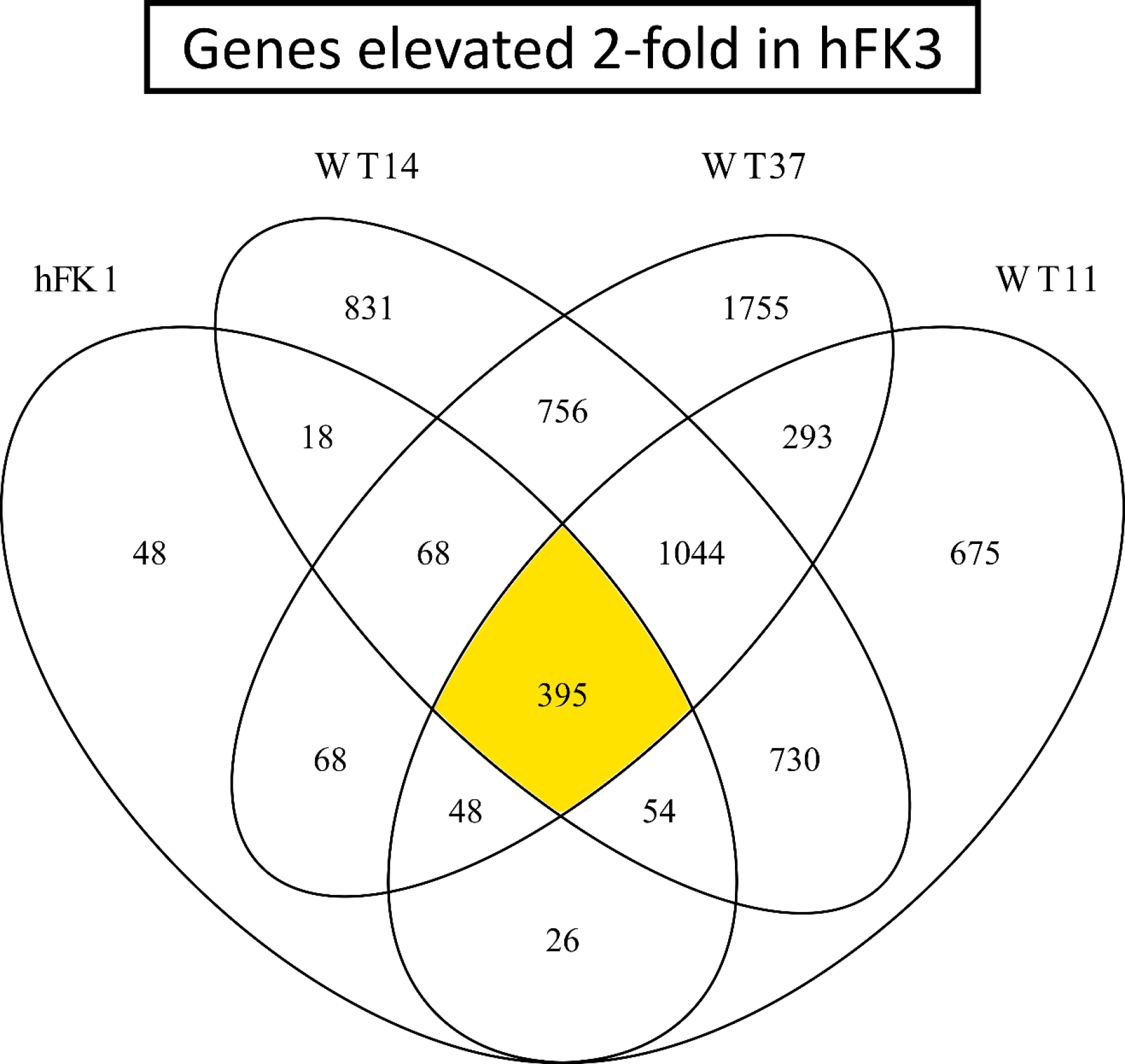
We selected a set of 395 genes that were found by intersecting all the gene sets that were upregulated at least 2-fold (log2foldChange > 1) in hFK3 (the mature fetal developmental fraction) with respect to hFK1, WT11, WT14, and WT37, and found that they are related to epithelial differentiation (Fig. 2D and Table S2). Shown is a Venn diagram of differentially expressed genes with log2foldChange > 1. Each oval represents a set of genes that were found to be upregulated in hFK3 with respect to the labeled cell fraction (hFK1, WT11, WT14, or WT37). We note that the enrichment that we observed was quite robust and did not depend much on the exact parameters for choosing these genes (e.g. the exact threshold for log2foldchange etc.)

**Figure S2:**
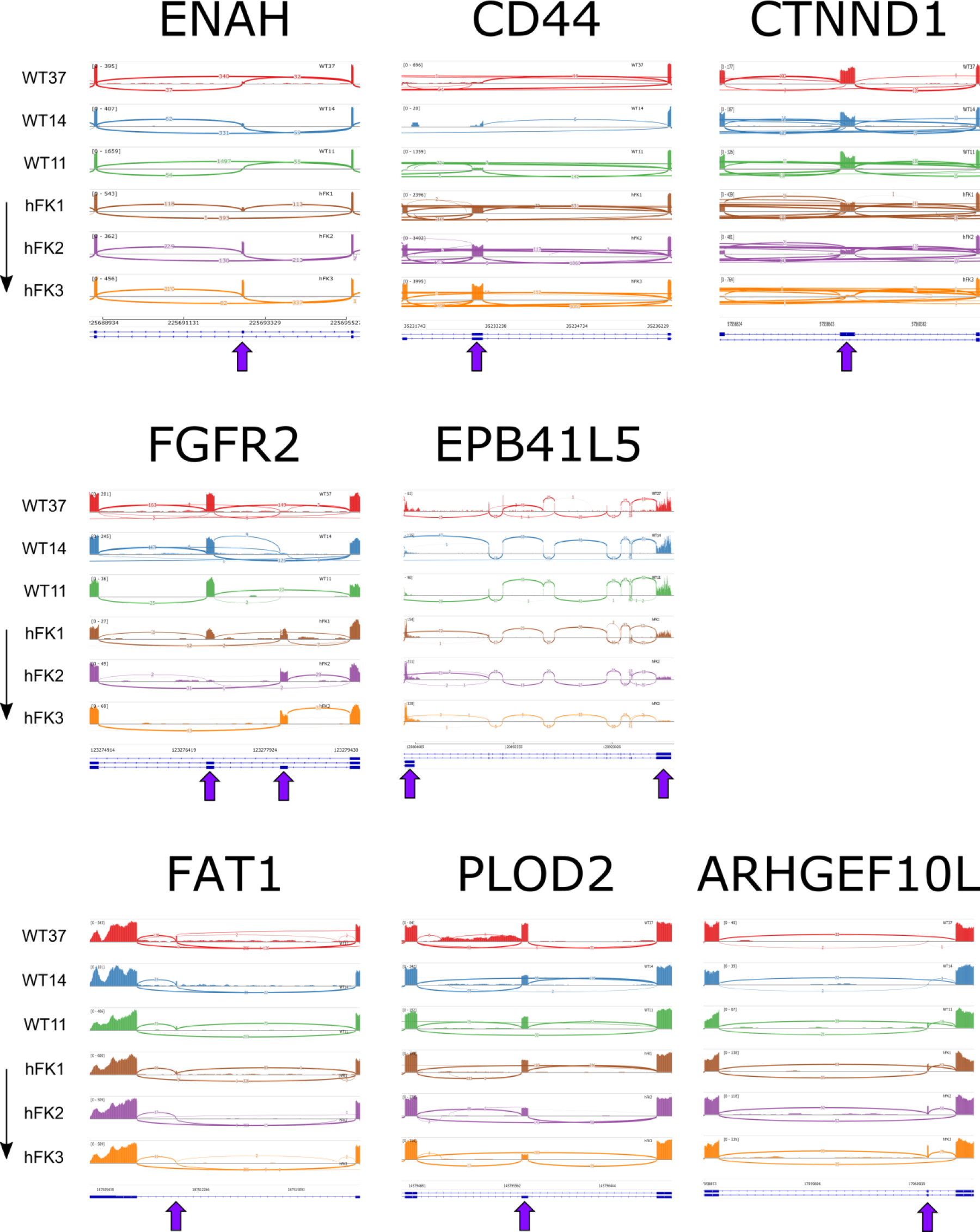
Cells at the early stage of human kidney development (hFK1) have a mesenchymal splice-isoform profile that is similar to that observed in blastemal-predominant Wilms’ tumor patient-derived xenografts (WT37, WT14, and WT11). Shown are Sashimi plots for the genes ENAH [1–3], CD44 [4,5], CTNND1 [3,6], FGFR2 [6,7], and EPB41L5 [3,8]. These genes were previously shown to be alternatively spliced in mesenchymal and epithelial tissues. The genes ENAH and CD44 contain cassette exons that are low in the Wilms’ tumor xenografts and increase during kidney development, whereas CTNND1 contains a cassette exon that is high in Wilms’ tumors and decreases during kidney development. FGFR2 contains two mutually exclusive exons, one of which is high in Wilms’ tumors and decreases during kidney development, and the other which is low in Wilms’ tumors and increases during kidney development. Likewise, EPB41L5 has two isoforms with alternative 3’ ends, one of which is predominant in the Wilms’ tumor xenografts and decreases during kidney development, and the other which is low in Wilms’ tumors and increases during kidney development. FAT1 [1,9] and PLOD2 [1,10,11], two genes for which alternative splicing was previously found to be regulated by the RNA binding protein RBFOX2 [1], contain cassette exons that are highly expressed in Wilms’ tumors and decrease during kidney development, while ARHGEF10L [12–14], a gene for which alternative splicing was previously found to be regulated by the RNA binding proteins ESRP1 and ESRP2 [12], contains a cassette exon that is low in Wilms’ tumors and increases during kidney development.

**Figure S3:**
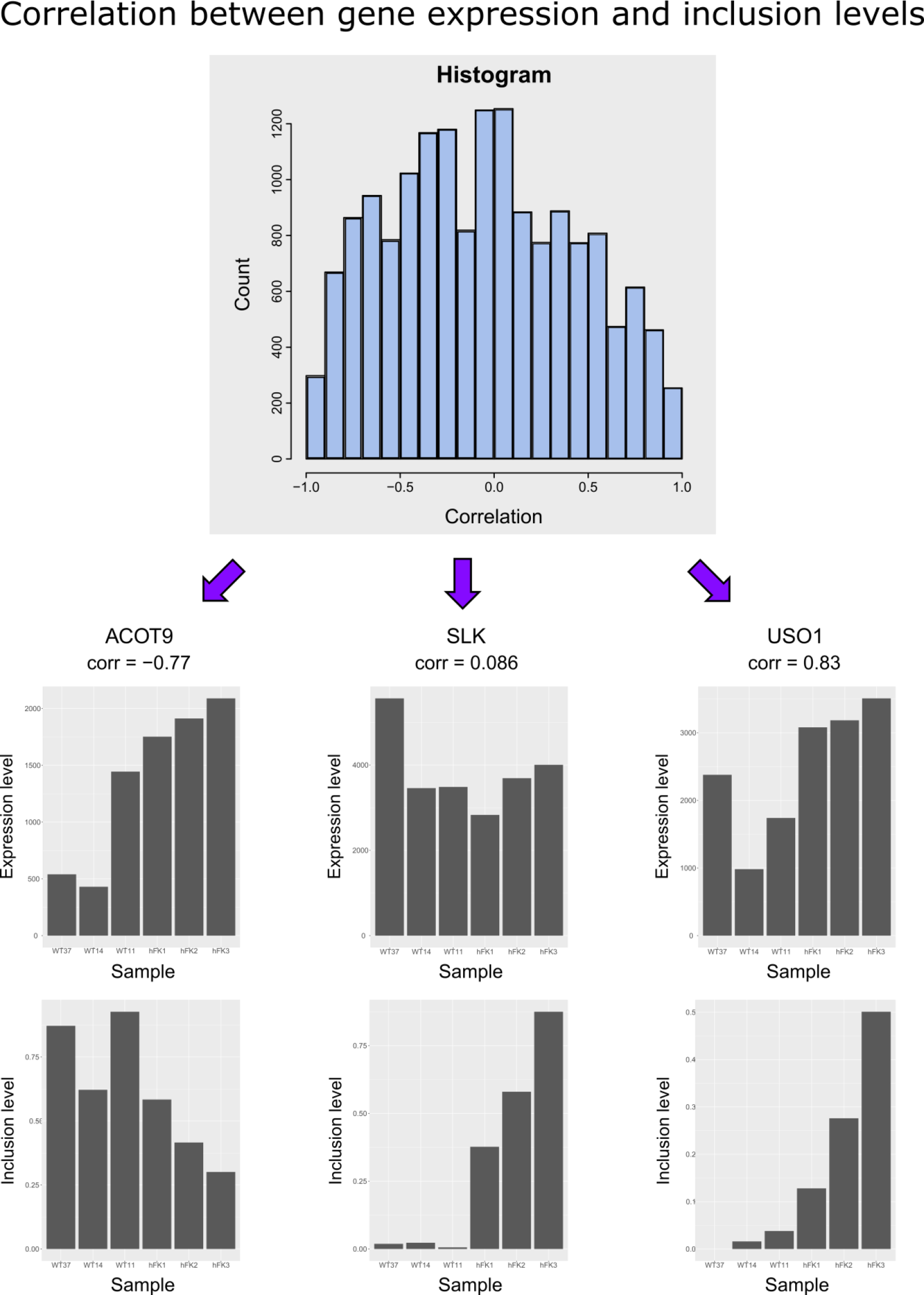
In some genes expression and splicing are strongly correlated or anti-correlated while in others they are not. Shown is a histogram of correlation values between expression levels and cassette exon inclusion levels within the same gene, as well as specific examples.

**Figure S4:**
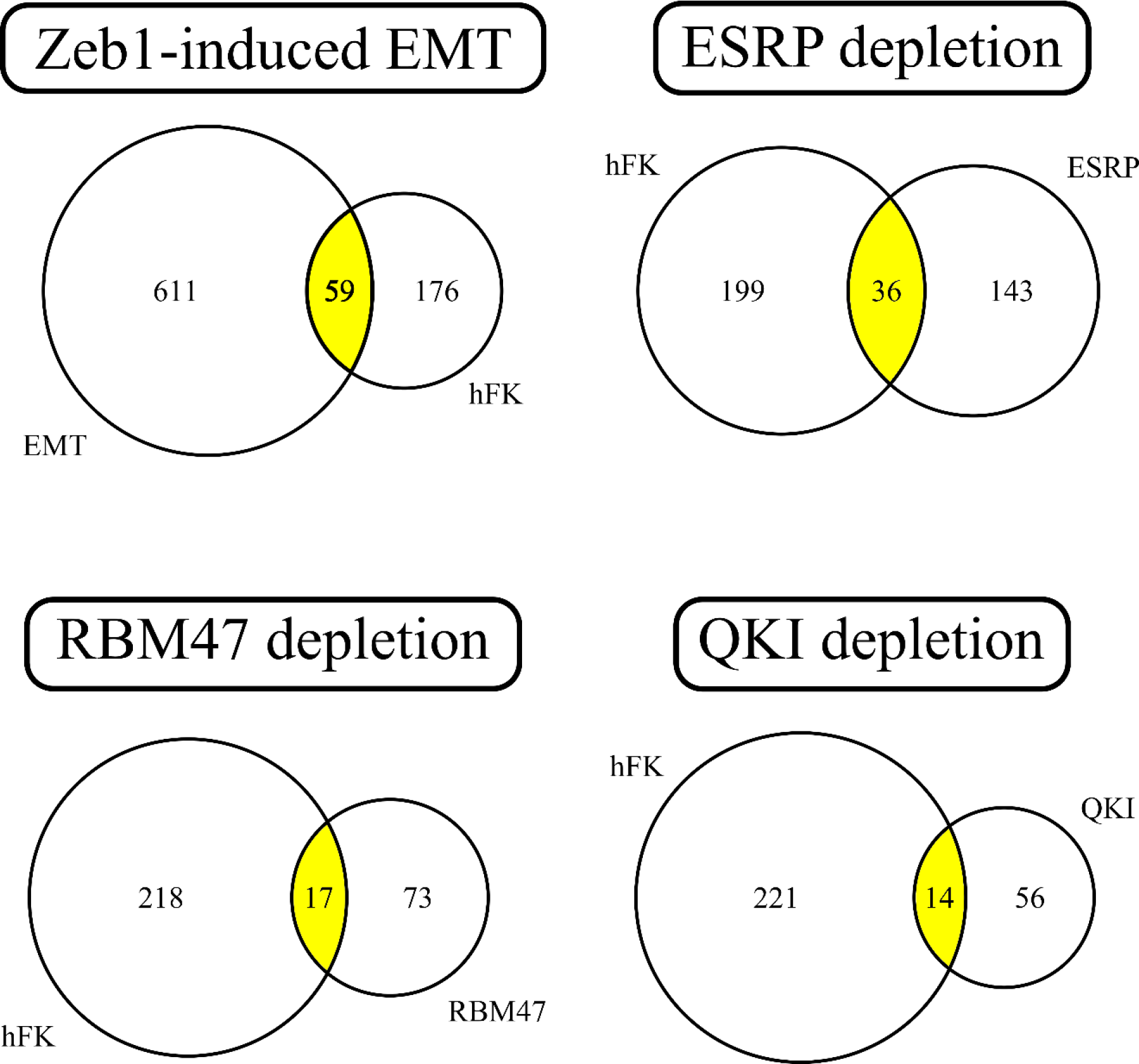
A comparison of cassette exons that were found in the present study to be alternatively spliced between cell fractions representing early (hFK1) and late (hFK3) stages of human kidney development vs. alternatively spliced cassette exons that were previously observed by Yang *et al.* [11] in cells from a human H358 epithelial non-small cell lung cancer (NSCLC) cell line undergoing Epithelial to Mesenchymal Transition (EMT). Shown are Venn diagrams comparing the findings of our study (“hFK”) to four different experiments performed in Yang *et al.* [11]: (1) Zeb1-induced EMT, (2) ESRP1/2 depletion, (3) RBM47 depletion, and (4) QKI depletion. In order to perform a consistent comparison, we followed the selection criteria of Yang *et al.* [11] and selected cassette exons for which the FDR < 0.05 and the absolute difference in inclusion levels > 0.05. For example, we found that the cassette exons located within the genes CD44 and ENAH are common to both our experiments (hFK) and also to the Zeb1-induced EMT, ESRP1/2 depletion, and RBM47 depletion experiments.

**Figure S5:**
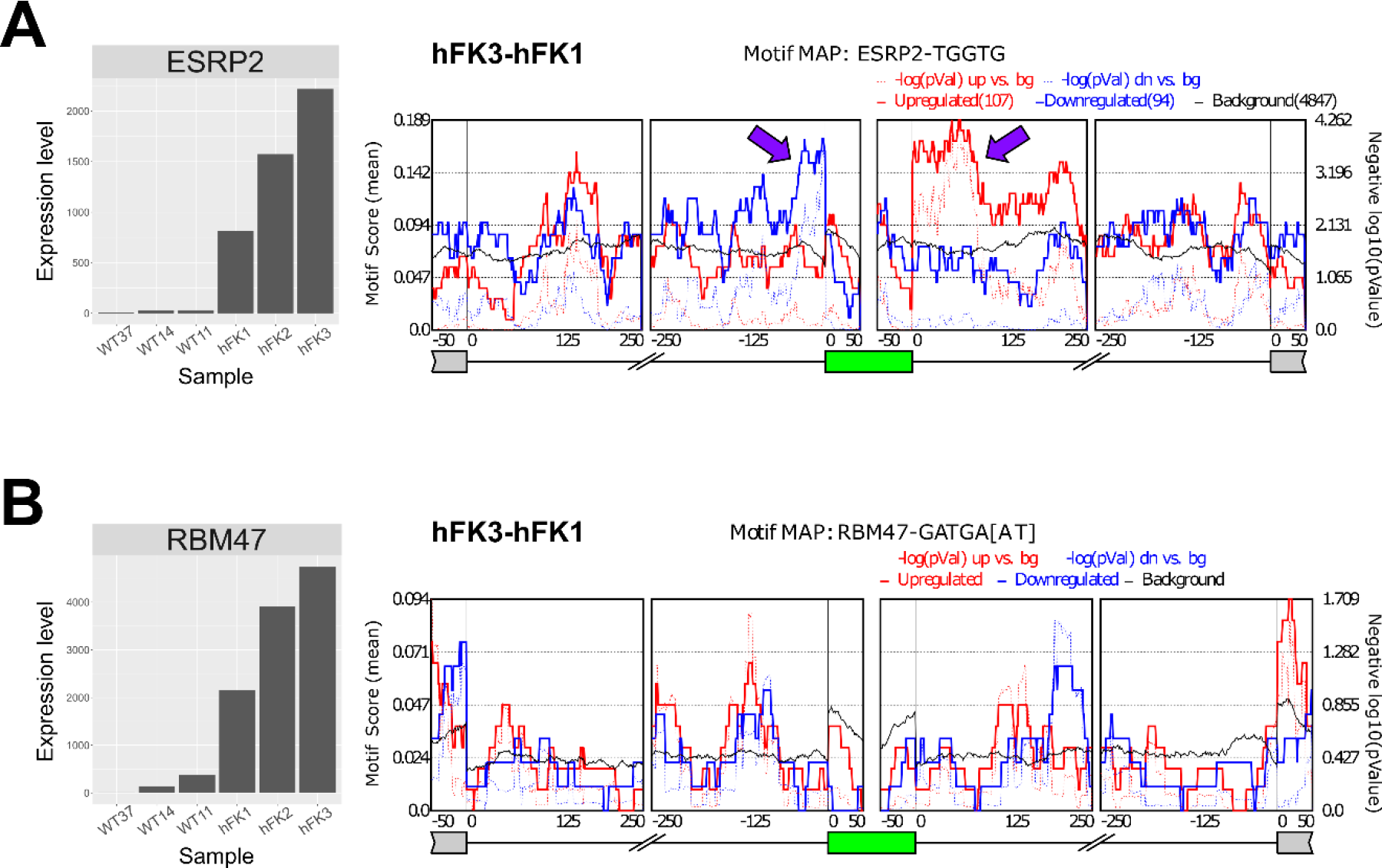
RNA binding motif enrichment analysis indicates that the mRNA binding protein ESRP2 regulates splice isoform switching during human kidney development. (A) ESRP2 expression levels are low in all Wilms’ tumor samples and monotonically increase along human kidney development, starting with moderate levels in hFK1 and reaching a maximum in hFK3. Likewise, exons that were enhanced in hFK3 (with respect to hFK1) are enriched for ESRP2 binding sites at their downstream 3’ flanking introns (red curve), while exons that are silenced in hFK3 are enriched for ESRP2 binding sites at their upstream 5’ flanking introns (blue curve). (B) Although the RNA binding protein RBM47 [11] is differentially over-expressed in hFK3 vs. hFK1, we did not find significant motif enrichment for this gene.

**Figure S6:**
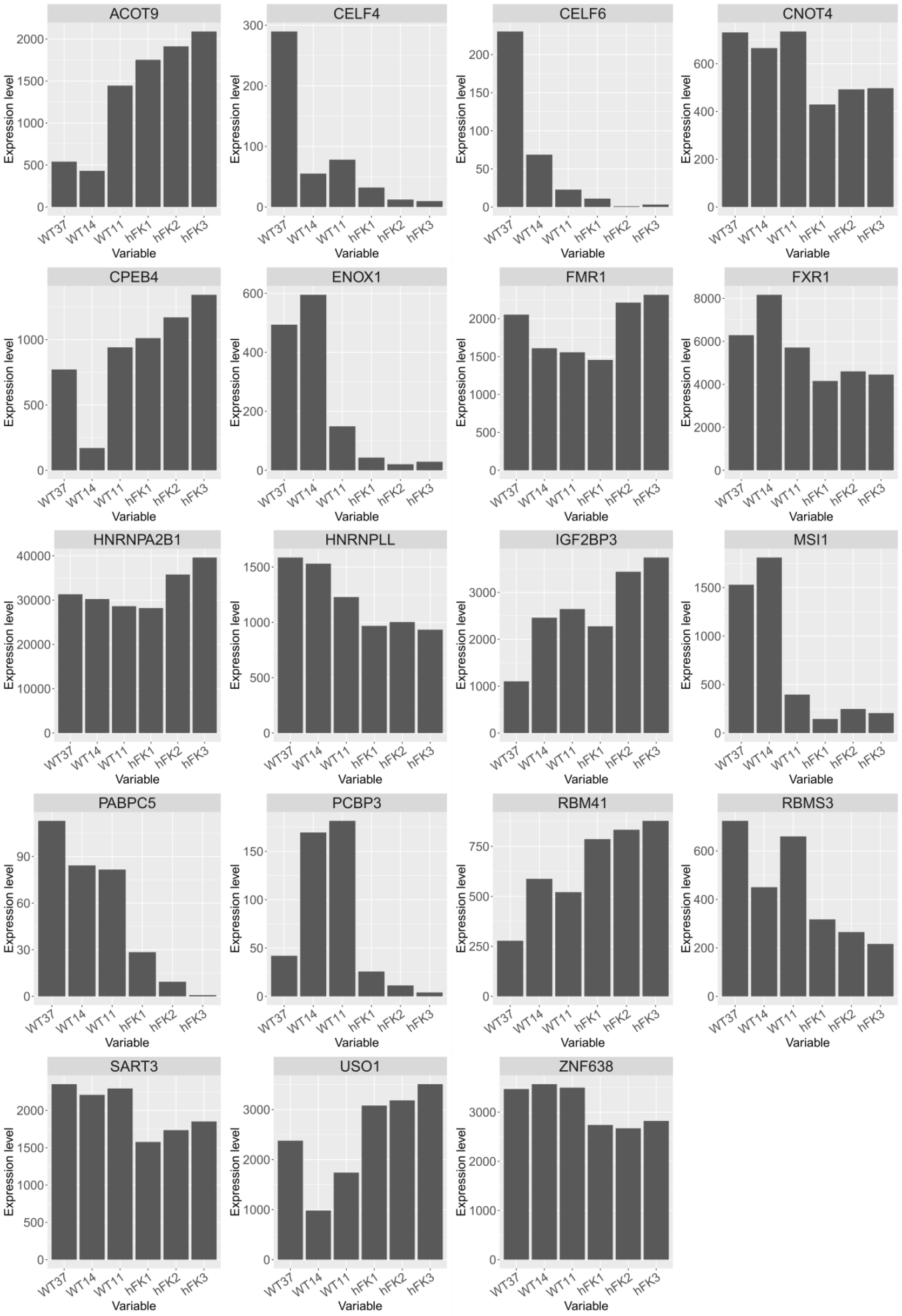
Additional RNA binding proteins that are differentially expressed between the cell fractions representing the early (hFK1) and late (hFK3) stages of human kidney development or between the blastemal-predominant Wilms’ tumor patient-derived xenografts (WT37, WT14, and WT11) and hFK3. These RBP’s might also be involved in splicing regulation in the developing fetal kidney or in the development of Wilms’ tumors.

## REFERENCES

1. Hohenstein P, Pritchard-Jones K, Charlton J. The yin and yang of kidney development and Wilms’ tumors. Genes Dev. 2015;29: 467–482. doi:10.1101/gad.256396.114

2. Little M, Georgas K, Pennisi D, Wilkinson L. Kidney Development: Two Tales of Tubulogenesis. In: Thornhill BA, Chevalier RL, editors. Current Topics in Developmental Biology. 2010. pp. 193–229. doi:10.1016/S0070-2153(10)90005-7

3. Gilbert SF. Developmental Biology. 6th edition. Sunderland (MA): Sinauer Associates; 2000. Intermediate Mesoderm. Available from: https://www.ncbi.nlm.nih.gov/books/NBK10089/.

4. Gilbert SF. Intermediate Mesoderm. Developmental Biology. Sunderland (MA): Sinauer Associates; 2000. Available: http://www.ncbi.nlm.nih.gov/books/NBK10089/

5. Davidoff AM. Wilms Tumor. Adv Pediatr. Elsevier Inc; 2012;59: 247–267. doi:10.1016/j.yapd.2012.04.001

6. Huff V. Wilms’ tumours: about tumour suppressor genes, an oncogene and a chameleon gene. Nat Rev Cancer. Nature Publishing Group; 2011;11: 111–121. doi:10.1038/nrc3002

7. Popov SD, Sebire NJ, Vujanic GM. Wilms’ Tumour – Histology and Differential Diagnosis. In: van den Heuvel-Eibrink MM, editor. Wilm Tumor. Brisbane, Australia; 2016. doi:10.15586/codon.wt.2016.ch1

8. Pode-Shakked N, Pleniceanu O, Gershon R, Shukrun R, Kanter I, Bucris E, et al. Dissecting Stages of Human Kidney Development and Tumorigenesis with Surface Markers Affords Simple Prospective Purification of Nephron Stem Cells. Sci Rep. 2016;6: 23562. doi:10.1038/srep23562

9. Shapiro IM, Cheng AW, Flytzanis NC, Balsamo M, Condeelis JS, Oktay MH, et al. An emt-driven alternative splicing program occurs in human breast cancer and modulates cellular phenotype. PLoS Genet. 2011;7. doi:10.1371/journal.pgen.1002218

10. Di Modugno F, Iapicca P, Boudreau A, Mottolese M, Terrenato I, Perracchio L, et al. Splicing program of human MENA produces a previously undescribed isoform associated with invasive, mesenchymal-like breast tumors. Proc Natl Acad Sci U S A. 2012;109: 19280–5. doi:10.1073/pnas.1214394109

11. Warzecha CC, Shen S, Xing Y, Carstens RP. The epithelial splicing factors ESRP1 and ESRP2 positively and negatively regulate diverse types of alternative splicing events. RNA Biol. 2009;6: 546–562. doi:10.4161/rna.6.5.9606

12. Sneath RJ, Mangham DC. The normal structure and function of CD44 and its role in neoplasia. Mol Pathol. 1998;51: 191–200. doi:10.1136/mp.51.4.191

13. Brown RL, Reinke LM, Damerow MS, Perez D, Chodosh L a., Yang J, et al. CD44 splice isoform switching in human and mouse epithelium is essential for epithelial-mesenchymal transition and breast cancer progression. J Clin Invest. 2011;121: 1064–1074. doi:10.1172/JCI44540

14. Pode-Shakked N, Gershon R, Tam G, Omer D, Gnatek Y, Kanter I, et al. Evidence of In Vitro Preservation of Human Nephrogenesis at the Single-Cell Level. Stem Cell Reports. ElsevierCompany.; 2017;9. doi:10.1016/j.stemcr.2017.04.026

15. Pode-Shakked N, Gershon R, Tam G, Omer D, Gnatek Y, Kanter I, et al. Evidence of In Vitro Preservation of Human Nephrogenesis at the Single-Cell Level. Stem Cell Reports. ElsevierCompany.; 2017;9. doi:10.1016/j.stemcr.2017.04.026

16. Metsuyanim S, Harari-Steinberg O, Buzhor E, Omer D, Pode-Shakked N, Ben-Hur H, et al. Expression of stem cell markers in the human fetal kidney. PLoS One. 2009;4. doi:10.1371/journal.pone.0006709

17. Pode-Shakked N, Shukrun R, Mark-Danieli M, Tsvetkov P, Bahar S, Pri-Chen S, et al. The isolation and characterization of renal cancer initiating cells from human Wilms’ tumour xenografts unveils new therapeutic targets. EMBO Mol Med. 2013;5: 18–37. doi:10.1002/emmm.201201516

18. Pode-Shakked N, Metsuyanim S, Rom-Gross E, Mor Y, Fridman E, Goldstein I, et al. Developmental tumourigenesis: NCAM as a putative marker for the malignant renal stem/progenitor cell population. J Cell Mol Med. 2009;13: 1792–808. Available: http://www.ncbi.nlm.nih.gov/pubmed/20187302

19. Brunskill EW, Aronow BJ, Georgas K, Rumballe B, Valerius MT, Aronow J, et al. Atlas of Gene Expression in the Developing Kidney at Microanatomic Resolution. Dev Cell. Elsevier Ltd; 2008;15: 781–791. doi:10.1016/j.devcel.2008.09.007

20. Magella B, Adam M, Potter AS, Venkatasubramanian M, Chetal K, Hay SB, et al. Cross-platform single cell analysis of kidney development shows stromal cells express Gdnf. Dev Biol. Elsevier Inc.; 2017;434: 1–12. doi:10.1016/j.ydbio.2017.11.006

21. Adam M, Potter AS, Potter SS. Psychrophilic proteases dramatically reduce single cell RNA-seq artifacts: A molecular atlas of kidney development. Development. 2017;1: dev.151142. doi:10.1242/dev.151142

22. Chen J, Bardes EE, Aronow BJ, Jegga AG. ToppGene Suite for gene list enrichment analysis and candidate gene prioritization. Nucleic Acids Res. 2009; doi:10.1093/nar/gkp427

23. Subramanian A, Tamayo P, Mootha VK, Mukherjee S, Ebert BL. Gene set enrichment analysis : A knowledge-based approach for interpreting genome-wide. Proc Natl Acad Sci U S A. 2005;102: 15545–15550. Available: http://www.broad.mit.edu/gsea/

24. Warzecha CC, Sato TK, Nabet B, Hogenesch JB, Carstens RP. ESRP1 and ESRP2 Are Epithelial Cell-Type-Specific Regulators of FGFR2 Splicing. Mol Cell. Elsevier Ltd; 2009;33: 591–601. doi:10.1016/j.molcel.2009.01.025

25. Hovhannisyan RH, Warzecha CC, Carstens RP. Characterization of sequences and mechanisms through which ISE/ISS-3 regulates FGFR2 splicing. Nucleic Acids Res. 2006;34: 373–385. doi:10.1093/nar/gkj407

26. Warzecha CC, Carstens RP. Complex changes in alternative pre-mRNA splicing play a central role in the epithelial-to-mesenchymal transition (EMT). Semin Cancer Biol. Elsevier Ltd; 2012;22: 417–427. doi:10.1016/j.semcancer.2012.04.003

27. Shen S, Park JW, Lu Z, Lin L, Henry MD, Wu YN, et al. rMATS: robust and flexible detection of differential alternative splicing from replicate RNA-Seq data. Proc Natl Acad Sci U S A. 2014;111: E5593–601. doi:10.1073/pnas.1419161111

28. Thorvaldsdóttir H, Robinson JT, Mesirov JP. Integrative Genomics Viewer (IGV): High-performance genomics data visualization and exploration. Brief Bioinform. 2013;14: 178–192. doi:10.1093/bib/bbs017

29. Bebee TW, Park JW, Sheridan KI, Warzecha CC, Cieply BW, Rohacek AM, et al. The splicing regulators Esrp1 and Esrp2 direct an epithelial splicing program essential for mammalian development. Elife. 2015;4: 1–27. doi:10.7554/eLife.08954

30. Wang Z, Burge CB. Splicing regulation: From a parts list of regulatory elements to an integrated splicing code. Rna. 2008;14: 802–813. doi:10.1261/rna.876308.802

31. Wang ET, Sandberg R, Luo S, Khrebtukova I, Zhang L, Mayr C, et al. Alternative isoform regulation in human tissue transcriptomes. Nature. 2008;456: 470–476. doi:10.1038/nature07509

32. Braeutigam C, Rago L, Rolke A, Waldmeier L, Christofori G, Winter J. The RNA-binding protein Rbfox2: An essential regulator of EMT-driven alternative splicing and a mediator of cellular invasion. Oncogene. Nature Publishing Group; 2014;33: 1082–1092. doi:10.1038/onc.2013.50

33. Yeowell HN, Walker LC. Tissue specificity of a new splice form of the human lysyl hydroxylase 2 gene. Matrix Biol. 1999;18: 179–187. doi:10.1016/S0945-053X(99)00013-X

34. Yang Y, Park JW, Bebee TW, Warzecha CC, Guo Y, Shang X, et al. Determination of a Comprehensive Alternative Splicing Regulatory Network and Combinatorial Regulation by Key Factors during the Epithelial-to-Mesenchymal Transition. Mol Cell Biol. 2016;36: 1704–1719. doi:10.1128/MCB.00019-16

35. Bangru S, Arif W, Seimetz J, Bhate A, Chen J, Rashan EH, et al. Alternative splicing rewires Hippo signaling pathway in hepatocytes to promote liver regeneration. Nat Struct Mol Biol. 2018;25: 928–939. doi:10.1038/s41594-018-0129-2

36. Bhate A, Parker DJ, Bebee TW, Ahn J, Arif W, Rashan EH, et al. ESRP2 controls an adult splicing programme in hepatocytes to support postnatal liver maturation. Nat Commun. Nature Publishing Group; 2015;6: 8768. doi:10.1038/ncomms9768

37. Park JW, Jung S, Rouchka EC, Tseng YT, Xing Y. rMAPS: RNA map analysis and plotting server for alternative exon regulation. Nucleic Acids Res. 2016;44: W333–W338. doi:10.1093/nar/gkw410

38. Ray D, Kazan H, Cook KB, Weirauch MT, Najafabadi HS, Li X, et al. A compendium of RNA-binding motifs for decoding gene regulation. Nature. Nature Publishing Group; 2013;499: 172–177. doi:10.1038/nature12311

39. Black DL, Burge CB, Anderson ES, Lin C-H, Xiao X, Stoilov P. The cardiotonic steroid digitoxin regulates alternative splicing through depletion of the splicing factors SRSF3 and TRA2B. Rna. 2012;17: 1041–1049. doi:10.1261/rna.032912.112

40. Dittmar KA, Jiang P, Park JW, Amirikian K, Wan J, Shen S, et al. Genome-Wide Determination of a Broad ESRP-Regulated Posttranscriptional Network by High-Throughput Sequencing. Mol Cell Biol. 2012;32: 1468–1482. doi:10.1128/MCB.06536-11

41. Jin Y, Suzuki H, Maegawa S, Endo H, Sugano S, Hashimoto K, et al. A vertebrate RNA-binding protein Fox-1 regulates tissue-specific splicing via the pentanucleotide GCAUG. EMBO J. 2003;22: 905–912. doi:10.1093/emboj/cdg089

42. Picelli S, Björklund AK, Reinius B, Sagasser S, Winberg G, Sandberg R. Tn5 transposase and tagmentation procedures for massively scaled sequencing projects. Genome Res. 2014;24: 2033–2040. doi:10.1101/gr.177881.114

43. Picelli S, Björklund ÅK, Faridani OR, Sagasser S, Winberg G, Sandberg R. Smart-seq2 for sensitive full-length transcriptome profiling in single cells. Nat Methods. 2013;10: 1096–8. doi:10.1038/nmeth.2639

44. Shalek AK, Satija R, Adiconis X, Gertner RS, Gaublomme JT, Raychowdhury R, et al. Single-cell transcriptomics reveals bimodality in expression and splicing in immune cells. Nature. 2013;498: 236–240. doi:10.1038/nature12172.Single-cell

45. Brunskill EW, Park J-S, Chung E, Chen F, Magella B, Potter SS. kh. Development. 2014;141: 3093–3101. doi:10.1242/dev.110601

46. Wineberg Y, Bar-Lev TH, Futorian A, Ben-Haim N, Armon L, Ickowicz D, et al. Single-cell RNA sequencing reveals mRNA splice isoform switching during kidney development. bioRxiv. Cold Spring Harbor Laboratory; 2019; doi:10.1101/688564

47. Warzecha CC, Jiang P, Amirikian K, Dittmar K a, Lu H, Shen S, et al. An ESRP-regulated splicing programme is abrogated during the epithelial-mesenchymal transition. EMBO J. Nature Publishing Group; 2010;29: 3286–3300. doi:10.1038/emboj.2010.195

48. Bebee TW, Sims-Lucas S, Park JW, Bushnell D, Cieply B, Xing Y, et al. Ablation of the epithelial-specific splicing factor Esrp1 results in ureteric branching defects and reduced nephron number. Dev Dyn. 2016;245: 991–1000. doi:10.1002/dvdy.24431

49. Kim D, Pertea G, Trapnell C, Pimentel H, Kelley R, Salzberg SL. TopHat2: accurate alignment of transcriptomes in the presence of insertions, deletions and gene fusions. Genome Biol. 2013;14: R36. doi:10.1186/gb-2013-14-4-r36

50. Anders S, Pyl PT, Huber W. HTSeq-A Python framework to work with high-throughput sequencing data. Bioinformatics. 2015;31: 166–169. doi:10.1093/bioinformatics/btu638

51. Love MI, Huber W, Anders S. Moderated estimation of fold change and dispersion for RNA-seq data with DESeq2. Genome Biol. 2014;15: 550. doi:10.1186/s13059-014-0550-8

52. Katz Y, Wang ET, Silterra J, Schwartz S, Wong B, Thorvaldsdóttir H, et al. Quantitative visualization of alternative exon expression from RNA-seq data. Bioinformatics. 2015;31: 2400–2402. doi:10.1093/bioinformatics/btv034

## REFERENCES

1. Shapiro IM, Cheng AW, Flytzanis NC, Balsamo M, Condeelis JS, Oktay MH, et al. An emt-driven alternative splicing program occurs in human breast cancer and modulates cellular phenotype. PLoS Genet. 2011;7. doi:10.1371/journal.pgen.1002218

2. Di Modugno F, Iapicca P, Boudreau A, Mottolese M, Terrenato I, Perracchio L, et al. Splicing program of human MENA produces a previously undescribed isoform associated with invasive, mesenchymal-like breast tumors. Proc Natl Acad Sci U S A. 2012;109: 19280–5. doi:10.1073/pnas.1214394109

3. Warzecha CC, Shen S, Xing Y, Carstens RP. The epithelial splicing factors ESRP1 and ESRP2 positively and negatively regulate diverse types of alternative splicing events. RNA Biol. 2009;6: 546–562. doi:10.4161/rna.6.5.9606

4. Sneath RJ, Mangham DC. The normal structure and function of CD44 and its role in neoplasia. Mol Pathol. 1998;51: 191–200. doi:10.1136/mp.51.4.191

5. Brown RL, Reinke LM, Damerow MS, Perez D, Chodosh L a., Yang J, et al. CD44 splice isoform switching in human and mouse epithelium is essential for epithelial-mesenchymal transition and breast cancer progression. J Clin Invest. 2011;121: 1064–1074. doi:10.1172/JCI44540

6. Warzecha CC, Sato TK, Nabet B, Hogenesch JB, Carstens RP. ESRP1 and ESRP2 Are Epithelial Cell-Type-Specific Regulators of FGFR2 Splicing. Mol Cell. Elsevier Ltd; 2009;33: 591–601. doi:10.1016/j.molcel.2009.01.025

7. Hovhannisyan RH, Warzecha CC, Carstens RP. Characterization of sequences and mechanisms through which ISE/ISS-3 regulates FGFR2 splicing. Nucleic Acids Res. 2006;34: 373–385. doi:10.1093/nar/gkj407

8. Warzecha CC, Carstens RP. Complex changes in alternative pre-mRNA splicing play a central role in the epithelial-to-mesenchymal transition (EMT). Semin Cancer Biol. Elsevier Ltd; 2012;22: 417–427. doi:10.1016/j.semcancer.2012.04.003

9. Braeutigam C, Rago L, Rolke A, Waldmeier L, Christofori G, Winter J. The RNA-binding protein Rbfox2: An essential regulator of EMT-driven alternative splicing and a mediator of cellular invasion. Oncogene. Nature Publishing Group; 2014;33: 1082–1092. doi:10.1038/onc.2013.50

10. Yeowell HN, Walker LC. Tissue specificity of a new splice form of the human lysyl hydroxylase 2 gene. Matrix Biol. 1999;18: 179–187. doi:10.1016/S0945-053X(99)00013-X

11. Yang Y, Park JW, Bebee TW, Warzecha CC, Guo Y, Shang X, et al. Determination of a Comprehensive Alternative Splicing Regulatory Network and Combinatorial Regulation by Key Factors during the Epithelial-to-Mesenchymal Transition. Mol Cell Biol. 2016;36: 1704–1719. doi:10.1128/MCB.00019-16

12. Bebee TW, Park JW, Sheridan KI, Warzecha CC, Cieply BW, Rohacek AM, et al. The splicing regulators Esrp1 and Esrp2 direct an epithelial splicing program essential for mammalian development. Elife. 2015;4: 1–27. doi:10.7554/eLife.08954

13. Bangru S, Arif W, Seimetz J, Bhate A, Chen J, Rashan EH, et al. Alternative splicing rewires Hippo signaling pathway in hepatocytes to promote liver regeneration. Nat Struct Mol Biol. 2018;25: 928–939. doi:10.1038/s41594-018-0129-2

14. Bhate A, Parker DJ, Bebee TW, Ahn J, Arif W, Rashan EH, et al. ESRP2 controls an adult splicing programme in hepatocytes to support postnatal liver maturation. Nat Commun. Nature Publishing Group; 2015;6: 8768. doi:10.1038/ncomms9768

